# A novel mutant allele uncouples brassinosteroid-dependent and independent functions of BRI1

**DOI:** 10.1101/605923

**Authors:** Eleonore Holzwart, Nina Glöckner, Herman Höfte, Klaus Harter, Sebastian Wolf

## Abstract

Plants depend on an array of cell surface receptors to integrate extracellular signals with developmental programs. One of the best-studied receptors is BRASSINOSTEROID INSENSITIVE 1 (BRI1), which upon binding of its hormone ligands forms a complex with shape-complimentary co-receptors and initiates a signal transduction cascade leading to a wide range of responses. BR biosynthetic and receptor mutants have similar growth defects on the macroscopic level, which had initially led to the assumption of a largely linear signalling pathway. However, recent evidence suggests that BR signalling is interconnected with a number of other pathways through a variety of different mechanisms. We recently described that feedback information from the cell wall is integrated at the level of the receptor complex through interaction with RLP44. Moreover, BRI1 is required for a second function of RLP44, the control of procambial cell fate. Here, we report on a *BRI1* mutant, *bri1*^*cnu4*^, which differentially affects canonical BR signalling and RLP44 function in the vasculature. While BR signalling is only mildly impaired, *bri1*^*cnu4*^ mutants show ectopic xylem in the position of procambium. Mechanistically, this is explained by an increased association of RLP44 and the mutated BRI1 protein, which prevents the former from acting in vascular cell fate maintenance. Consistent with this, the mild BR response phenotype of *bri1*^*cnu4*^ is a recessive trait, whereas the RLP44-mediated xylem phenotype is semi-dominant. Our results highlight the complexity of plant plasma membrane receptor function and provide a tool to dissect BR signalling-related roles of BRI1 from its non-canonical functions.

**One sentence summary:** A novel mutant allows to dissect brassinosteroid signalling related and non-canonical functions of the receptor-like kinase BRI1.

## Introduction

Plant cells perceive a multitude of extracellular signals through a battery of plasma membrane-bound receptors that are crucial for the integration of environmental and developmental signals. The response to the growth-regulatory brassinosteroid (BR) phytohormones is mediated by one of the best-characterized plant signalling pathways (Singh and Savaldi-Goldstein, 2015) initiated by a receptor complex containing the leucine-rich repeat receptor-like kinase BRASSINOSTEROID INSENSITIVE 1 (BRI1) (Li and Chory, 1997) and its co-receptors of the SOMATIC EMBRYOGENESIS RECEPTOR-LIKE KINASE (SERK) family (Ma *et al.*, 2016, Hohmann *et al.*, 2017). Binding of the brassinosteroid ligand mediates hetero-dimerization of BRI1 and SERK family members such as BRI1-ASSOCIATED RECEPTOR KINASE 1 (BAK1) (Li *et al.*, 2002, Nam and Li, 2002), which in turn triggers extensive auto- and trans-phosphorylation of the intracellular BAK1 and BRI1 kinase domains (Hohmann *et al.*, 2017). The activated kinases recruit and activate downstream BR signalling components, which eventually leads to vast changes in gene expression mediated by BR signalling-regulated transcription factors such as BRASSINAZOLE-RESISTANT 1 (BZR1) (Wang *et al.*, 2002) and BRI1-EMS-SUPPRESSOR 1 (BES1) (Yin *et al.*, 2002). Among the transcriptional targets of these transcription factors, cell wall related genes are strongly overrepresented, consistent with a growth-regulatory function of BR signalling (Sun *et al.*, 2010, Yu *et al.*, 2011, Chaiwanon and Wang, 2015). Recently, we reported that the state of the cell wall is connected to BR signalling through a feedback mechanism mediated by the RECEPTOR-LIKE PROTEIN 44 (RLP44). Plants in which the activity of the important cell wall modifying enzyme PECTIN METHYLESTERASE (PME) is impaired through ectopic expression of a PME inhibitor protein (PMEIox), BR signalling is activated in a compensatory response that includes transcriptional upregulation of PMEs to prevent cell (wall) rupture (Wolf *et al.*, 2012). RLP44 is sufficient to activate BR signalling, likely by acting as a scaffold to promote association of BRI1 and BAK1 (Wolf *et al.*, 2014), and this interaction is not affected by increasing BR levels. Thus, information from the cell wall is integrated with BR signalling activity at the level of the plasma membrane. Furthermore, RLP44 is under transcriptional control of BRI1 and is able to promote activity of a second LRR-RLK complex, containing the receptor for the phytosulfokine (PSK) peptide, PSK RECEPTOR 1 (PSKR1), through the same scaffolding mechanism as observed for the activation of BR signalling (Holzwart *et al.*, 2018). As a result, both BRI1 and RLP44 are required for full functionality of PSK signalling in the vasculature, demonstrated by the observation that *bri1* null mutants, *rlp44* mutants, and PSK-related mutants share the same vascular phenotype in the primary Arabidopsis root: ectopic xylem cells in the position of the procambium (Holzwart *et al.*, 2018). Interestingly, hypomorphic mutants of BRI1 with intermediate growth phenotypes and BR biosynthetic mutants with strong growth phenotypes show wild type-like xylem, suggesting that BRI1,s role in BR signalling is independent from its role in procambial maintenance. Here, we further dissect BRI1 function through the characterization of a novel *bri1* allele, which is only marginally affected in canonical BR signalling, but strongly affected in RLP44-mediated control of procambial cell fate. These observations demonstrate that the function of BRI1 in BR signalling can be uncoupled from its emerging additional functions.

## Results

### Two novel suppressor mutants of PMEIox

We have previously described that plants overexpressing a pectin methylesterase inhibitor protein (PMEIox) show a pleiotropic growth phenotype caused by cell wall feedback signalling. We have used these plants to perform a genetic screen which identified the *comfortably numb (cnu) 1* and *cnu2* suppressor mutants affected in the BR receptor BRI1 (Wolf *et al.*, 2012) and RLP44 (Wolf *et al.*, 2014), respectively. Reduced pectin methylesterase activity in PMEIox leads to a compensatory upregulation of BR signalling, which restores cell wall integrity but causes directional growth phenotypes as a secondary effect (Wolf *et al.*, 2012). RLP44 is required and sufficient for enhancing BR signalling in response to cell wall modification (Wolf *et al.*, 2014), presumably by promoting the interaction between BRI1 and its co-receptor BAK1 (Holzwart *et al.*, 2018). From the *cnu* suppressor screen we identified two new extragenic suppressor mutants, which we called *cnu3* and *cnu4* (Fig. 1A). Similar to *cnu1* and *cnu2*, both *cnu3* and *cnu4* strongly suppressed the macroscopic PMEIox growth phenotype in seedlings, with the exception of a residual root waving phenotype in *cnu3*, as indicated by measurement of the vertical growth index (Grabov *et al.*, 2005) (vertical distance between hypocotyl junction and root tip divided by root length) (Fig 1A). As adult plants, *cnu3* and *cnu4* appeared similar to wild type plants, in contrast to their parental line PMEIox (Fig. 1B). Moreover, *cnu3* and *cnu4* showed suppression of the malformed and short silique phenotype of PMEIox (Fig. 1C). Quantitative real time PCR analysis revealed that transcript levels of the BR signalling marker gene *DWF4* in *cnu3* and *cnu4* is intermediate between Col-0 and PMEIox, suggesting partial suppression of PMEIox-mediated activation of BR signalling (Fig 1D). Consistent with this notion, and similar to the *cnu1* (mutated in *BRI1*) and *cnu2* (mutated in *RLP44*) suppressor mutants, *cnu3* and *cnu4* were more resistant than Col-0 to the depletion of endogenous BR by propiconazole (PPZ) (Hartwig *et al.*, 2012), but showed a relatively normal response to exogenous application of epi-brassinolide (BL), in contrast to the largely insensitive *cnu1* mutant (Fig. 1E).

**Figure 1.**
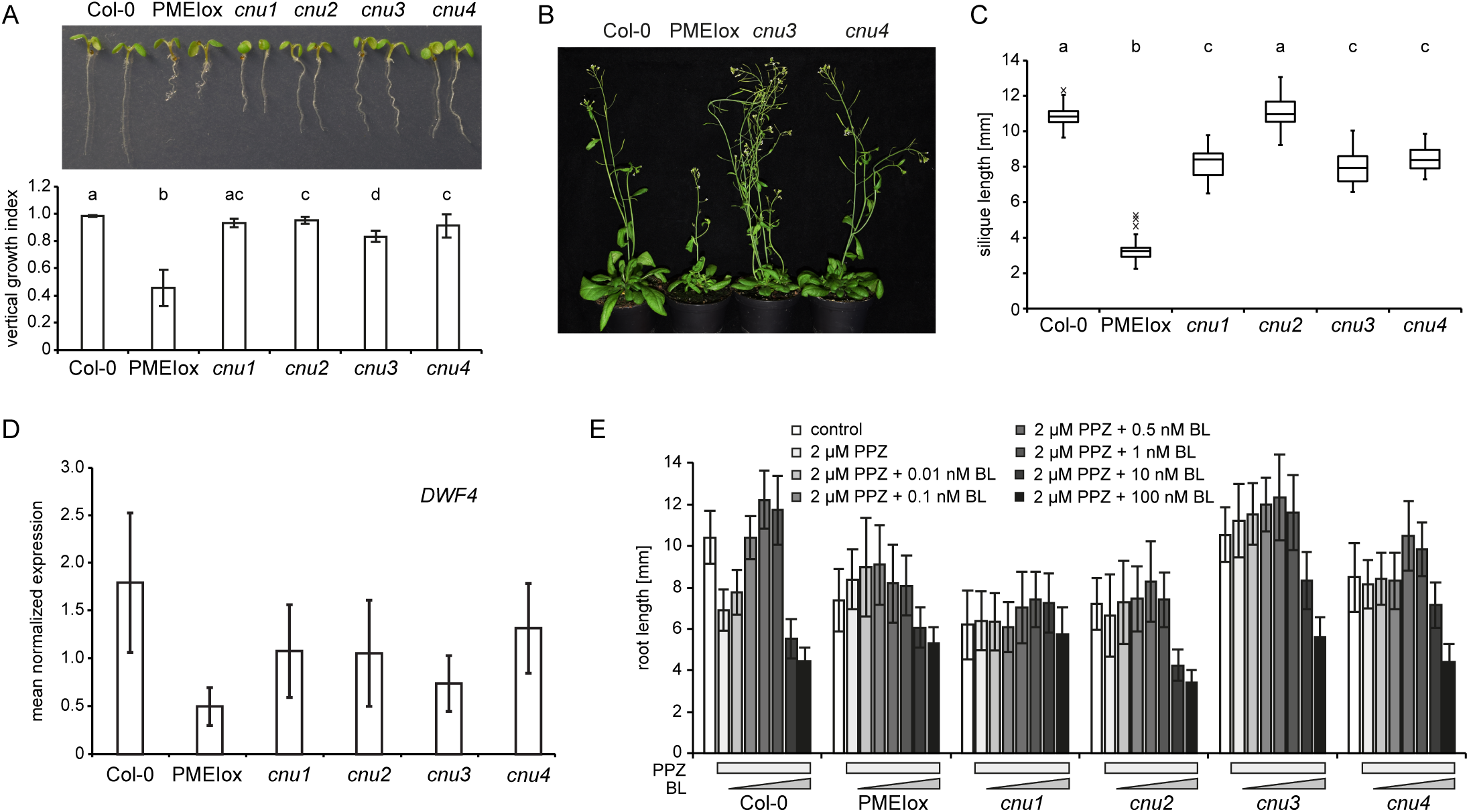
Identification of the PMEIox suppressor mutants *cnu3* and *cnu4*. **A**, seedling morphology and root vertical growth index (Grabov *et al.*, 2005) of Col-0, PMEIox, the previously published PMEIox suppressor mutants *cnu1* (Wolf *et al.*, 2012) and *cnu2* (Wolf *et al.*, 2014), and the two novel suppressor mutants, *cnu3* and *cnu4*. Letters indicate statistically significant difference according to one-way ANOVA with p < 0.05 (n = 13). **B**, Adult *cnu3* and *cnu4* mutants resemble wild type plants, in contrast to the PMEIox parental line. **C**, Silique length of Col-0, PMEIox, and the four PMEIox suppressor mutants *cnu1* to *cnu4*. Box plots in (A) indicate interquartile range (box), median (bar) and 1.5x IQR (whiskers), outliers are indicated with a cross, n = 24-36. **D**, qRT-PCR analysis of the BR biosynthetic gene *DWF4* in wild type (Col-0), PMEIox and the *cnu1* to *cnu4* the suppressor mutants. Bars depict average ± S.D., n = 3. E, Root length response of Col-0, PMEIox and the *cnu1* to *cnu4* suppressor mutants to BR depletion by PPZ and exogenous application of BL. Bars depict average ± S.D., n = 19-53.

### The *cnu3* and *cnu4* suppressor mutants carry two novel hypomorphic alleles of *bri1*

To gain insight into the relationship of *cnu3, cnu4*, and the previously described *rlp44* mutant *cnu2*, we performed allelism tests by crossing the different suppressor mutants with each other. F1 plants resulting from a cross between *cnu3* and *cnu4* showed suppression of PMEIox growth defects (Supplemental Fig. S1), whereas F1 plants generated by crossing with *cnu2* showed the PMEIox phenotype (Supplemental Fig. S1). This suggests that *cnu3* and *cnu4* are mutated in the same gene, which is, however, different from *RLP44*. As we had previously identified a PMEIox suppressor mutation in the BR receptor, we sequenced *BRI1* in the novel mutants. We revealed a mutation in *cnu3* leading to exchange of arginine 769, located in the extracellular membrane-proximal region, to tryptophan (R769W). In *cnu4*, we detected a SNP leading to the exchange of glycine 746, located in the last LRR repeat of the extracellular domain, to serine (G746S) (Fig. 2A). To test whether these variants were causative for the PMEIox suppressor phenotype, we complemented *cnu3* and *cnu4* by expressing GFP-tagged BRI1 under the control of its native 5, regulatory sequences. Transgenic BRI1-GFP expression resulted in restoration of the PMEIox phenotype or even a dwarf phenotype (Supplemental Fig. S2A), presumably because expression of BRI1 in these hypomorphic mutants in the presence of PMEIox-induced cell wall alterations can lead to excessive BRI1 activity detrimental to growth. Consistent with this assumption, our complementation lines were infertile and reminiscent of plants derived from a cross between PMEIox and BRI1 overexpressing plants (Friedrichsen *et al.*, 2000), which also showed extreme dwarfism and were unable to reproduce (Wolf *et al.*, 2012). To characterize the effect of the mutations in the absence of PMEIox-induced cell wall alterations, we crossed *cnu3* and *cnu4* to the Col-0 wild type, and genotyped the F2 population to identify individuals that contained the homozygous *bri1* mutations but had lost the PMEIox transgene. We called those mutants derived from *cnu3* and *cnu4 bri1*^*cnu3*^ and *bri1*^*cnu4*^, respectively. In sharp contrast to our previously identified PMEIox-suppressing mutant *bri1*^*cnu1*^ (Wolf *et al.*, 2012), both mutants showed relatively normal growth and were not strongly deviating from the wild type with respect to classical BR signalling hallmarks such as fertility, leaf shape, leaf colour, silique length, and marker gene expression (Fig. 2B-D). To assess the capacity of the *bri1* mutants to respond to altered BR levels, we grew seedlings on plates under BR-depleting conditions and externally applied varying concentration of BL. Depletion of BRs by PPZ reduced root length of 5-days-old seedlings to approximately 5 mm in all genotypes. Co-treatment with 0.5 nM BL completely restored Col-0 root length, whereas 1 nM of BL was required to achieve maximal root length in *bri1*^*cnu3*^ and *bri1*^*cnu4*^ (Fig. 2E). Further increase of BL led to growth depression in WT, and, to slightly lesser degree, in the *bri1*^*cnu3*^ and *bri1*^*cnu4*^ mutants. Thus, in accordance with the subtle growth phenotype, *bri1*^*cnu3*^ and *bri1*^*cnu4*^ were only mildly affected in their response to altered levels of BRs. In contrast, *bri1*^*cnu1*^ was much less responsive to exogenous BR and did not reach growth depression with the concentrations tested here (up to 10 nM) (Fig. 2E), as reported for other *bri1* hypomorphic alleles of similar strength (Sun *et al.*, 2017). Consistent with the mild growth phenotype, transformation with constructs encoding the two *BRI1* mutant versions alone or a combination of both mutations rescued hypomorphic *bri1-301* and *bri1-null* mutants (Fig 3A, B). The subcellular localization of *pBRI1*-expressed BRI1cnu4-GFP was indistinguishable from *pBRI1*-expressed BRI1-GFP (Fig 3C). Taken together, *bri1*^*cnu3*^ and *bri1*^*cnu4*^ are two weak BRI1 mutants with a mild growth phenotype.

**Figure 2.**
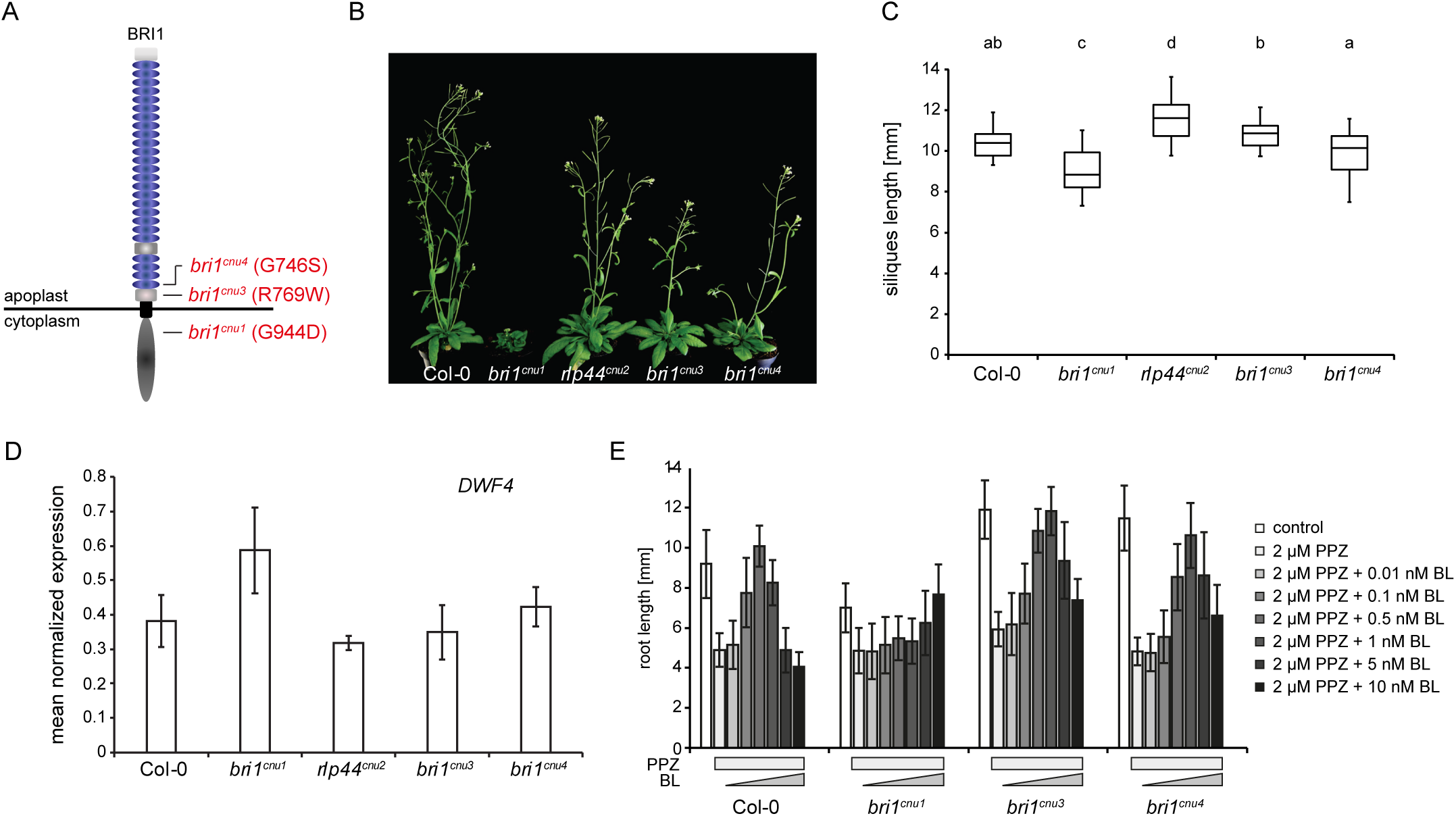
The *cnu3* and *cnu4* mutants are two novel alleles of *BRI1*. **A**, Schematic view of BRI1 with indicated position and amino acid substitution of the mutations in *bri1*^*cnu1*^, *bri1*^*cnu3*^ (derived from *cnu3*, but in the absence of the PMEIox transgene), and *bri1*^*cnu4*^ (derived from *cnu4*, but in the absence of the PMEIox transgene). **B**, Comparison of adult plant phenotype of Col-0, *bri1*^*cnu1*^, *rlp44*^*cnu2*^, *bri1*^*cnu3*^, and *bri1*^*cnu4*^. **C**, Silique length of Col-0, and the mutants derived from the *cnu1* to *cnu4* suppressor mutants. Box plots indicate interquartile range (box), median (bar) and 1.5x IQR (whiskers), n = 25-36. **D**, qRT-PCR analysis of the BR biosynthetic gene *DWF4* in wild type (Col-0), *bri1*^*cnu1*^, *rlp44*^*cnu2*^, *bri1*^*cnu3*^, and *bri1*^*cnu4*^. Bars depict average ± S.D., n = 3. **E**, Root length response of wild type (Col-0), *bri1*^*cnu1*^, *bri1*^*cnu3*^ and *bri1*^*cnu4*^ to BR depletion by PPZ and exogenous application of BL. Bars depict average ± S.D., n = 34-70.

**Figure 3.**
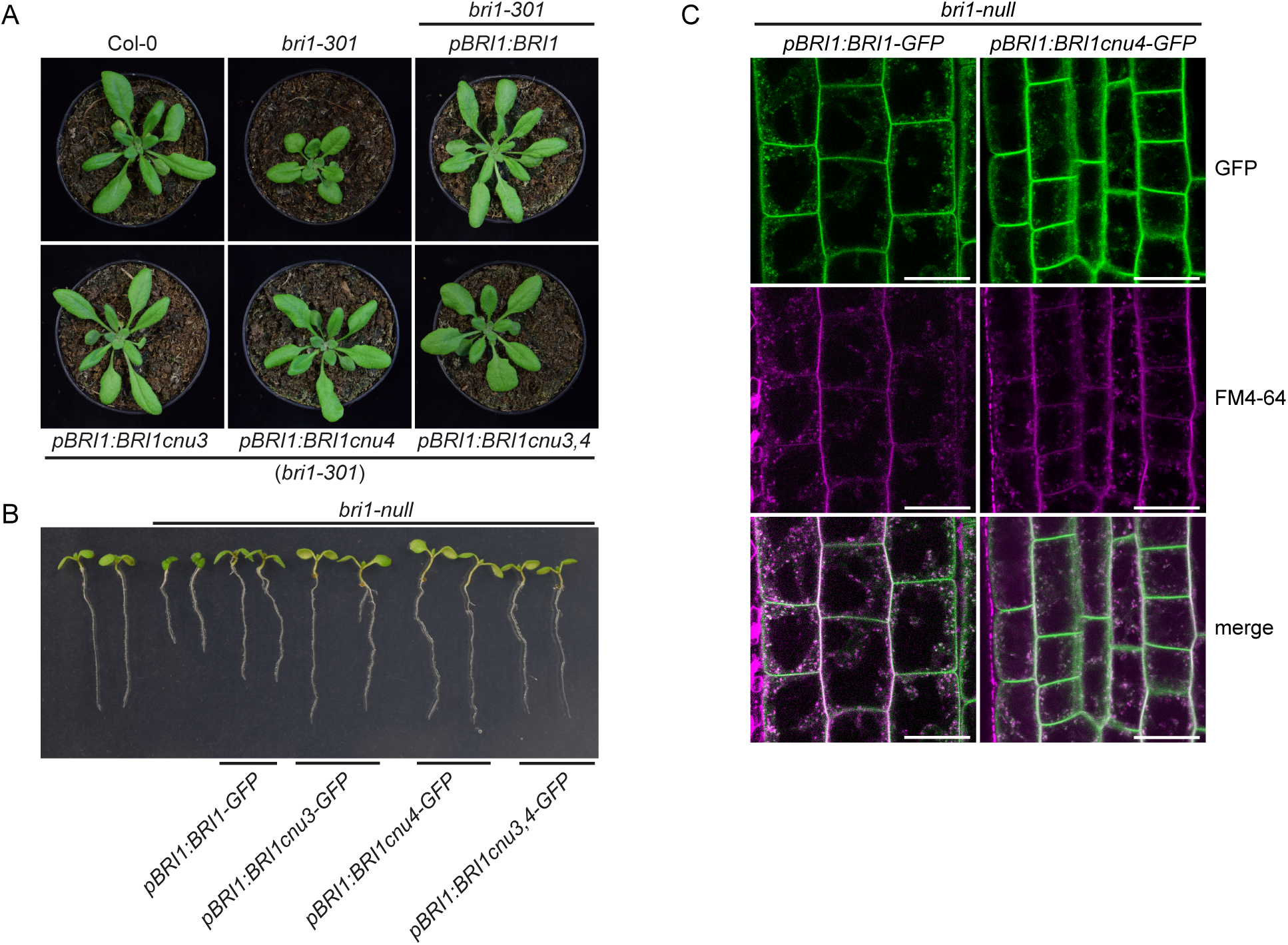
BRI1cnu4 and BRI1cnu3 proteins are functional. **A**, Mutant BRI1 constructs complement the hypomorphic *bri1-301* mutant. **B**, Constructs encoding mutated BRI1 versions complement the *bri-null* mutant. **C**, GFP fluorescence in root meristems of *bri1 null* mutants complemented with GFP fusion proteins from either the construct pBRI1:BRI1-GFP or pBRI1:BRI1cnu4-GFP, shows no apparent difference in subcellular localization. FM4-64 was used as an endocytic membrane tracer dye. Scale bars = 10 µm.

We have previously reported that *bri1* null but not *bri1* hypomorphic mutants show ectopic xylem cells in place of procambium in the *Arabidopsis* primary root. BRI1 controls vascular cell fate through a non-canonical, BR signalling-independent pathway acting through RLP44 and PSK signalling (Holzwart *et al.*, 2018). We therefore tested the xylem phenotype in *bri1*^*cnu4*^, expecting it would behave like other *bri1* hypomoprphic mutants such as *bri1*^*cnu1*^, *bri1-301*, and *bri1-5* (Noguchi *et al.*, 1999, Xu *et al.*, 2008, Wolf *et al.*, 2012, Holzwart *et al.*, 2018). In contrast, *bri1*^*cnu4*^ showed a strong increase in xylem cell number, comparable with *rlp44* mutants and slightly less pronounced than in *bri1*-null mutants (Fig. 4A) (Holzwart *et al.*, 2018). This clearly distinguishes *bri1*^*cnu4*^ from other BR-related mutants and suggests that the mutation in the BRI1cnu4 protein has a negative effect on RLP44 function. We reasoned that this could provide valuable insight into the mechanism of xylem cell fate determination by BRI1 and RLP44, concentrating on *bri1*^*cnu4*^ for the remainder of the study. We tested genetic interaction between *bri1*^*cnu4*^ and *rlp44*^*cnu2*^ by generating the double mutant and assessing its xylem phenotype. Simultaneous mutation of *rlp44* did not further enhance the *bri1*^*cnu4*^ mutant phenotype, suggesting that *bri1*^*cnu4*^ and *rlp44*^*cnu2*^ are affected in the same pathway with respect to xylem cell fate (Fig. 4A) Likewise, the subtle growth phenotype of *rlp44*^*cnu2*^ and *bri1*^*cnu4*^ was not aggravated in the double mutant (Fig. 4B).

**Figure 4.**
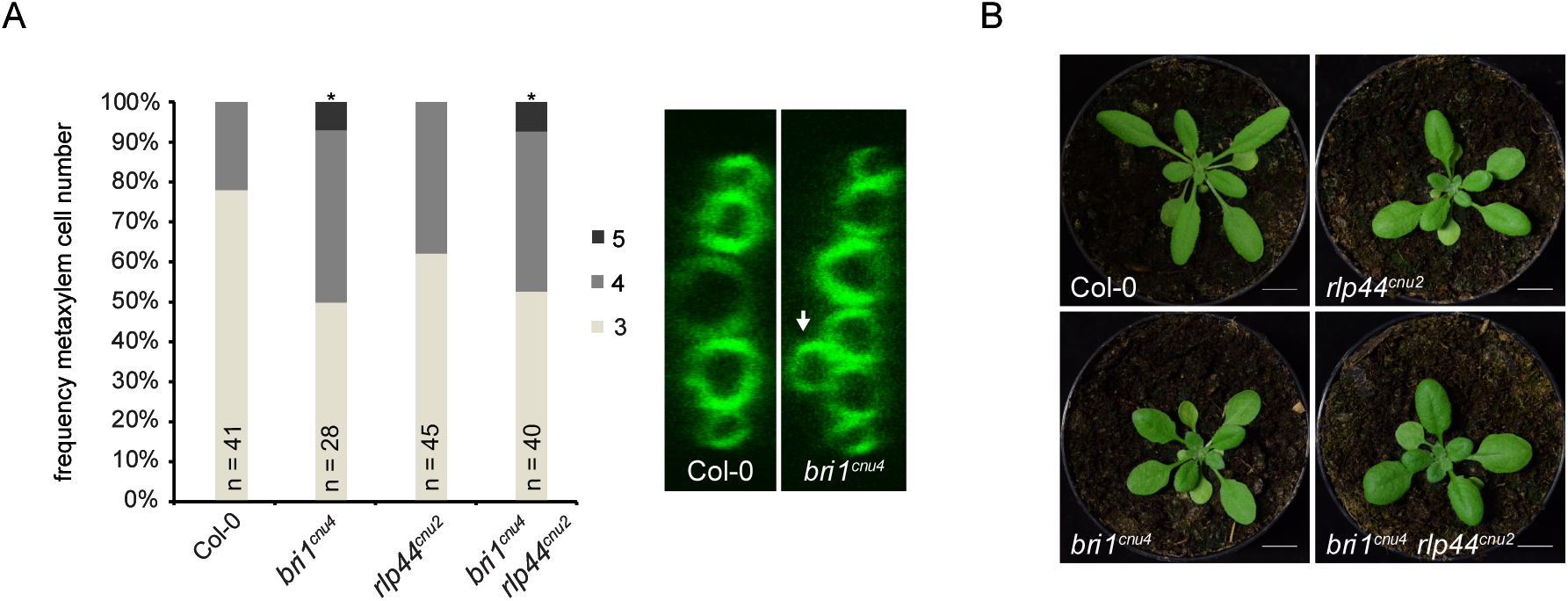
The mutation in *bri1*^*cnu4*^ negatively affects RLP44 function. **A**, Frequency of roots the indicated number of metaxylem cells in Col-0, *rlp44*^*cnu2*^, *bri1*^*cnu4*^, and the *rlp44*^*cnu2*^ *bri1*^*cnu4*^ double mutant. **B**, Morphological phenotype of Col-0, *rlp44*^*cnu2*^, *bri1*^*cnu4*^, and the *4*^*cnu2*^ *bri1*^*cnu4*^ double mutant.

### The *bri1*^*cnu4*^ mutant uncouples BRI1 roles in BR signalling and RLP-mediated control of cell fate

To further test our hypothesis that BRI1cnu4 negatively affects the function of RLP44 we assessed whether the mutation had a dominant effect. We analysed F1 hybrid seedlings derived from a cross of *bri1*^*cnu4*^ and Col-0, and revealed that the subtle BR insensitivity observed in *bri1*^*cnu4*^ root growth is a recessive trait (Fig. 5A). In line with this, the morphological phenotype of the F1 hybrids appeared closer to the wild type than to that of plants homozygous for the *bri1*^*cnu4*^ mutation (Fig. 5B). In addition, plants heterozygous for the *bri1*^*cnu4*^ mutation were not able to suppress PMEIox-mediated activation of BR signalling (Fig. 5C), indicating that *bri1*^*cnu4*^ rescues PMEIox in the *cnu4* mutant through reduced BR signalling strength. Intriguingly, despite the recessive nature of its BR signalling defect, the xylem phenotype of *bri1*^*cnu4*^ was clearly dominant in the F1 seedlings, supporting the idea that the mutation might directly or indirectly impair RLP44 function (Fig. 5D). Consistent with this hypothesis, expression of the *BRI1cnu4* transgene in the *bri1-301* hypomorphic mutant recapitulated the *bri1*^*cnu4*^ phenotype, whereas expression of wild type *BRI1* did not (Fig. 5E). Interestingly, RLP44-mediated activation of BR signalling was not blocked in *bri1*^*cnu4*^, as the phenotype of plants overexpressing *RLP44* in the *bri1*^*cnu4*^ background was intermediate between the overexpressing line and the mutant (Fig. 5F). This is in contrast to what was observed with overexpression of RLP44 in *bri1-null* (Holzwart *et al.*, 2018) and *bri1*^*cnu1*^, which harbours a mutation in the kinase domain (Wolf et al., 2014). Moreover, increasing the amount of RLP44 through transgenic expression under control of its own promoter rescued the mild BR response phenotype of *bri1*^*cnu4*^ (Supplemental Fig. S3), and partially rescued xylem cell number (Fig. 5G).

**Figure 5.**
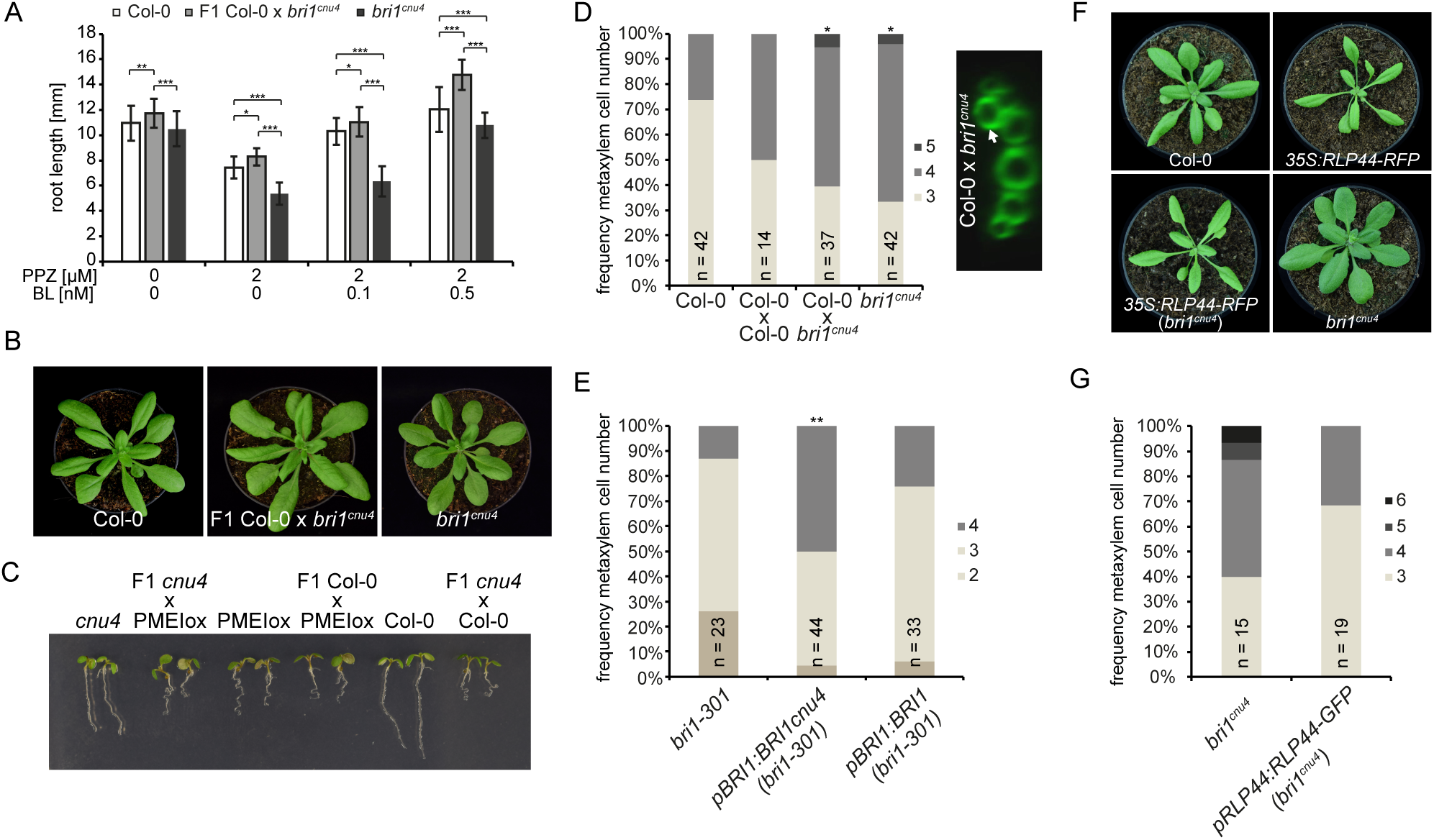
The *bri1*^*cnu4*^ mutant interferes with RLP44 function. **A**, Root length of 5-d-old F1 hybrid seedlings of a cross between *bri1*^*cnu4*^ and Col-0 after PPZ treatment and exogenous supply of BL. Bars indicate mean root length of 5-d-old seedlings ± SD (n = 22-49). Asterisks indicate significance with *p < 0.05, **p < 0.01, and ***p < 0.001 as determined by Tukey,s test after two-way ANOVA. Note that significance is only indicated for comparisons within each treatment. **B**, Morphological phenotype of Col-0, *bri1*^*cnu4*^, and F1 hybrid plants resulting from crossing the two genotypes. **C**, Suppression of PMEI5 overexpression phenotype (PMEIox) by the *bri1*^*cnu4*^ allele (*cnu4*) is a recessive trait, as indicated by the PMEIox-like phenotype of F1 plants from a cross between *cnu4* and Col-0. **D**, Ectopic xylem phenotype in *bri1*^*cnu4*^ and F1 plants from a cross between *bri1*^*cnu4*^ and Col-0. **E**, Expression of BRI1cnu4, but not of wildtype BRI1 in the *bri1-301* mutant results in supernumerary xylem cells. **F**, RLP44 overexpression can partially rescue the morphological phenotype of *bri1*^*cnu4*^. **G**, Increased expression of RLP44 can alleviate the *bri1*^*cnu4*^ phenotype. Asterisks indicate statistically significant difference from Col-0 based on Dunn,s post-hoc test with Benjamini-Hochberg correction after Kruskal-Wallis modified U-test (*p < 0.05).

To understand the mechanism by which BRI1cnu4 negatively affects RLP44 function, we analysed protein-protein interaction. To this end, we compared the association of RLP44 with BRI1 and BRI1cnu4 by immunoprecipitating *RLP44-RFP* in the Col-0 and *bri1*^*cnu4*^ background, respectively. Interestingly, BRI1cnu4 showed increased abundance in RLP44-containing complexes (Fig. 6A). Furthermore, split-ubiquitin assays in yeast supported stronger direct interaction between BRI1cnu4 and RLP44 as well as between BRI1cnu4 and BAK1 compared to wild type BRI1 (Fig. 6B). Thus, we assume that BRI1cnu4 exerts its effect on the maintenance of xylem cell fate by RLP44 sequestration thereby preventing RLP44 from acting in PSK signalling (Fig 6C).

**Figure 6.**
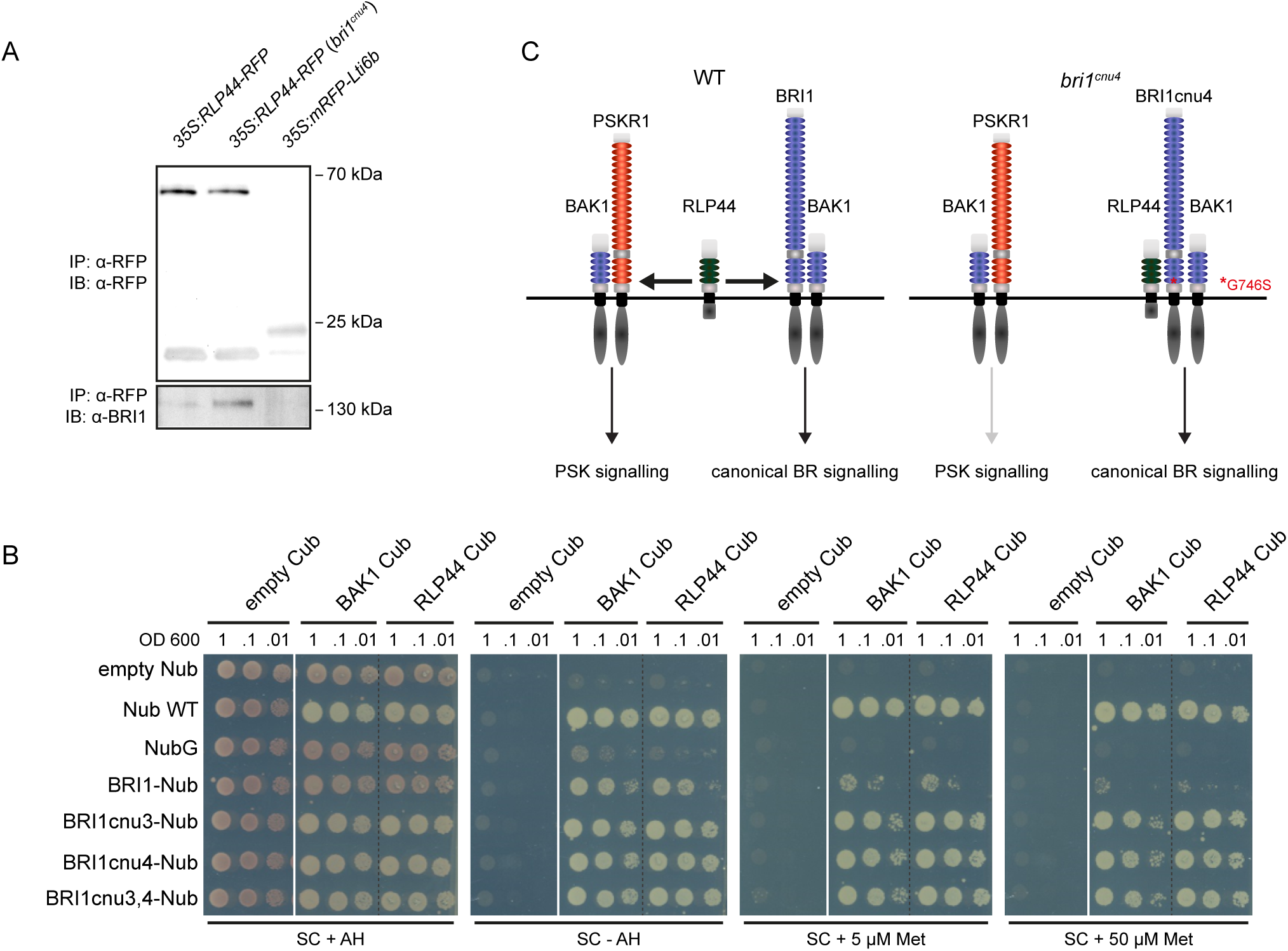
The BRI1cnu4 protein shows increased interaction with RLP44 and BAK1. **A**, Co-munoprecipitation of BRI1-GFP by RLP44-RFP from crude extracts of wild type (Col-0) and *bri1*^*cnu4*^ mutant plants. **B**, Mating-based split ubiquitin assays in yeast displaying the raction of BRI1, BRI1cnu3, BRI1cnu4 and BAK1 with RLP44. **C**, Model of RLP44 ractions with BRI1 and PSKR1 in the wild type and the *bri1*^*cnu4*^ mutant. The mutation at base of BRI1’s extracellular domain sequesters RLP44 and prevents it from promoting K/PSKR1 signalling.

## Discussion

We have previously shown that BRI1 have functions that are independent of classical BR signalling outputs mediated by the canonical BR signalling pathway (Holzwart et al., 2018). Here, we demonstrate that BRI1 mutants, depending on the nature of the allele, differentially affect these functions and can thus serve as a tool to uncouple canonical BR signalling-mediated from non-canonical effects. We isolated a novel *bri1* allele, *bri1*^*cnu4*^, and compared its impact on classical BR read-outs and the role of BRI1 in the maintenance of procambial cell fate, which depends on RLP44-mediated activation of PSK signalling (Holzwart *et al.*, 2018). These analyses revealed that BR signalling dependent BRI1 functions are only mildly affected in *bri1*^*cnu4*^, whereas we observed a strong negative effect on RLP44 function in the regulation of vascular cell fate. Interestingly, the same mutation we report here as *bri1*^*cnu4*^, G746A (G2236A on nucleic acid level) has been recently described as *bri1-711* in a tilling approach to obtain new *bri1* mutants (Sun *et al.*, 2017). Consistent with our results, *bri1-711* showed subtle growth defects and mild insensitivity to exogenous application of BL. In addition, the accumulation of non-phosphorylated BES1 as a readout of BR signalling was similar to that of the Col-0 WT in response to BL (Sun *et al.*, 2017). In contrast to our results obtained with *bri1*^*cnu4*^, other *bri1* hypomorphic mutants such as *bri1-301* and *bri1-5* have negligible effects on xylem cell fate in the root, despite their pronounced effect on BR signalling (Holzwart *et al.*, 2018). A possible explanation for the divergent effect of *bri1*^*cnu4*^ is provided by the observation that the BRI1cnu4 protein interacts more strongly with RLP44 than with wild type BRI1, and that additional RLP44 alleviates the *bri1*^*cnu4*^ xylem phenotype. From these observations we propose that BRI1cnu4 may sequester RLP44, which consequentially has a negative effect on PSK signalling. It has to be noted that in yeast mating-based split-ubiquitin system, BRI1cnu4 also shows increased interaction with its co-receptor BAK1, corroborating the complexity of receptor associations in the plasma membrane and the challenges associated with deciphering the multi-lateral interactions observed with many members of the LRR-RLK family (Stegmann *et al.*, 2017, Smakowska-Luzan *et al.*, 2018).

As revealed by the RLP44 interaction pattern, signalling integration and ramification is realised at the level of the receptor complex in the plasma membrane. Additional examples are the interaction of the BRI1-BAK1 complex with G-proteins to mediate sugar-responsive growth (Peng *et al.*, 2018), with the proton pumps of the P-ATPase type to regulate plasma membrane hyperpolarisation and wall swelling that precede cell elongation growth (Caesar *et al.*, 2011) and with the BAK1-interacting receptor-like kinase 3 (BIR3) that represses the activity of the complex in the absence of BR (Großeholz *et al.*, Imkampe *et al.*, 2017, Hohmann *et al.*, 2018)(Imkampe et al., 2017; Hohmann et al., 2018; Großeholz et al., 2019). In addition, BRI1 phosphorylates a homolog of the mammalian TGF-β receptor interacting protein/eIF3 eukaryotic translation initiation factor subunit TRIP-1 (Ehsan *et al.*, 2005). While the function of the latter protein is not completely clear at this stage, it seems at least conceivable that it bypasses the canonical BR signalling pathway, even if the morphological defects observed in plants expressing *TRIP-1* antisense RNA are reminiscent of BR-deficiency phenotypes (Jiang and Clouse, 2001).

The challenges emerging from the recent discoveries on the example of BRI1 is to understand of how distinct responses to extrinsic cues can be generated by the multifaceted network of a receptor in the plasma membrane. Thus, more sophisticated *in vivo* cell biological approaches in combination with genetic and biochemical tools are required to dissect and understand the function of this important signalling integrator, BRI1.

## Material and Methods

### Plant Material and growth conditions

All mutants and transgenic lines used in this study are in the Col-0 background. The *bri1*^*cnu1*^, *rlp44*^*cnu2*^, *bri1-null*, and *bri1-301* mutants have been described before (Xu *et al.*, 2008, Wolf *et al.*, 2012, Wolf *et al.*, 2014). The 35S:RLP44-RFP and pRLP44:RLP44-GFP (Wolf *et al.*, 2014, Holzwart *et al.*, 2018) described previously were used for crossing. All plants were grown in half-strength Murashige and Skoog (MS) medium supplemented with 1 % sucrose and 0.9 % plant agar. PPZ and 24-epi-brassinolide were added to the sterilized medium where appropriate.

### Plasmid generation

For mating-based split-ubiquitin assay (mbSUS) (Grefen et al., 2009), the coding sequence of RLP44, BAK1 and BRI1 in pDONR207 (Wolf et al., 2014) and were cloned into pMetYC-Dest. For generating the BRI1cnu3 Nub construct, primers BRI1_attB1_L + BRI1_attB2_R were used with gDNA of *bri1*^*cnu3*^ plants to create the full-length BRI1cnu3 cDNA in pDONR207. For generating the BRI1cnu4 Nub construct, primers BRI1_attB1_L + BRI1_attB2_R were used with gDNA of *bri1*^*cnu4*^ plants to create the full-length BRI1cnu4 cDNA in pDONR207. All other constructs used in this study were generated with GreenGate cloning as previously described (Lampropoulos et al., 2013). For generating BRI1cnu3,4 Nub construct, primers BRI1_attB1_L + BRI1_attB2_R wree used with the C-Module of BRI1cnu3,4 as a template to create the full-length BRI1cnu3,4 cDNA in pDONR207. The pDONR207 entry modules were recombined with pXNUbA22-Dest. For details regarding primers and constructs please see Supplemental Tables S1 and S2. For BRI1 (at4g39400) CDS GreenGate Cloning, three internal BsaI/Eco31I recognition sites were silently mutagenized via the generation of 4 PCR fragments with the primers BRI1_GGC_1F, BRI1_GGC_1R, BRI1_GGC_2F BRI1_GGC_2R, BRI1_GGC_3F, BRI1_GGC_3R, BRI1_GGC_4F and BRI1_GGC_4R as previously described (Holzwart et al., 2018). For generating the BRI1cnu4 module the second fragment was amplified with BRI1_GGC_2F BRI1_GGC_2R using gDNA of *bri1*^*cnu4*^ plants as template. For generating the BRI1cnu3 module the second fragment was amplified with BRI1_GGC_3F BRI1_GGC_3R with gDNA of *bri1*^*cnu3*^ plants. For the combined BRI1-cnu3,4 construtct, fragments 1 and 4 from BRI1 WT were combined with the second fragment of BRI1cnu4 and the third fragment of BRI1cnu3. PCR products of all combinations were gel purified, digested with Eco31I, subsequently ligated and processed according to the GreenGate protocol (Lampropoulos et al., 2013).

### Genotyping

Genotyping of *bri1*^*cnu1*^, *rlp44*^*cnu2*^, *bri1-301*, and *bri1-null* was described previously (Wolf et al., 2012, Wolf et al., 2014, Holzwart et al., 2018). For genotyping of the two new *bri1* alleles, we generated CAPS marker using primers bri1cnu3_CAPS_F, bri1cnu3_CAPS_R and restriction enzyme *Cfr*42I (*bri1*^*cnu3*^) or primers bri1cnu4_CAPS_F, bri1cnu4_CAPS_R with restriction enzyme *Bse*LI (*bri1*^*cnu4*^).

### Mating-based split ubiquitin assays

Yeast-based mbSUS assays were performed as described (Grefen *et al.*, 2009, Wolf *et al.*, 2014).

### Co-Immunoprecipitation

Material from transgenic plants expressing 35S:RLP44-RFP was frozen in liwuid nitrogen and ground to a fine powder using mortar and pestle. Extraction buffer (100 mM Tris-HCl (pH 8.0), 150 mM NaCl, 10% (v/v) Glycerol, 5 mM EDTA (Sigma-Aldrich), 2% (v/v) Igepal CA-630 (Sigma-Aldrich), 5 mM DTT (Sigma-Aldrich, added immediately prior to use), 1% (v/v) Protease Inhibitor Cocktail (Bimake, added immediately prior to use) was added to the frozen powder (2 ml per g fresh weight) and the homogenate was centrifuged at 12 000 x g and 4 °C after thawing. The supernatant was incubated with 15 µl of RFP-trap slurry (Chromotek) for 2 hours at 4°C on a rotary shaker. The beads were subsequently washed with extraction buffer 4 times and then boiled in 60 µl 2x SDS-PAGE sample buffer at 95 °C for 5 min. SDS-PAGE, Western Blotting and Immunological detection of RLP44-GFP and BRI1 was performed as described (Holzwart *et al.*, 2018).

### Confocal microscopy

GFP, FM4-64, and basic fuchsin fluorescence was analysed on a Leica SP5 microscope system equipped with a 63x water immersion objective using laser lines of 488 nm (GFP), 514 nm (basic fuchsin), and 543 nm (FM4-64). Fluorescence was recorded between 490 and 525 nm for GFP, between 530 and 600 nm for basic fuchsin, and between 600 nm and 720 nm for FM4-64. Images were analysed with Fiji.

### Xylem imaging

Basic fuchsin staining of seedling roots was performed as described (Holzwart *et al.*, 2018).

### Quantitative Real-Time PCR

Total RNA was extracted from 100 mg of tissue harvested form 5 day old seedlings using the GeneMATRIX Universal RNA Purification Kit (EURx/Roboklon). AMV Reverse Transcriptase Native according the manufacturer,s protocol (Roboklon E1372) with RiboLock Rnase Inhibitor (Thermo Fisher Scientific EO0381) was used for generating cDNA. PCR reactions were performed in a Rotor Gene Q 2plex cycler (Qiagen) using 1:40 diluted cDNA template, JumpStart Taq DNA polymerase (Sigma-Aldrich) and SYBR-GreenI (Sigma-Aldrich). Expression of DWF4 was normalized against at5g46630 (see Supplemetary Table S1 for oligonucleotide sequences).

## Author contribution

E.H. and N.G. performed experiments. E.H., N.G., and S.W. analysed data. H.H., K.H. and S.W. conceived and supervised research. S.W. agrees to serve as the author responsible for contact and ensures communication.

## Acknowledgements

The authors would like to thank Michael Hothorn for antiserum against BRI1 and Karin Schumacher for antiserum against RFP. Research in our laboratories was supported by the German Research Foundation (DFG) with grants WO 1660/6-1 (to S.W.), HA2146/22-1 (to K.H.), and CRC 1101-D02 (to K.H.). S.W. acknowledges funding by the DFG through the Emmy Noether Programme grant WO 1660/2. H.H. was financed in part by ANR grant “Pectosign”. The authors benefited from the IJPB Plant Observatory facilities. The IJPB benefits from the support of the LabEx Saclay Plant Sciences-SPS (ANR-10-LABX-0040-SPS).

---

**Supplemental Figure 1.** *cnu3* and *cnu4* are allelic mutants. PMEIox silique morphology (upper panel) and plant stature (lower panel) remain suppressed in F1 plants of a cross between *cnu3* and *cnu4*, whereas F1 plants of a cross between *cnu2* (carrying a mutation in RLP44) and *cnu4* show PMEIox phenotype.

**Supplemental Figure 2.** Mutant BRI1 constructs complement the *cnu3* and *cnu4* mutant**s**.

**Supplemental Figure 3.** RLP44 promotes BR response in the *bri1*^*cnu4*^ mutant. Response of Col-0, pRLP44:RLP44-GFP, *bri1*^*cnu4*^, and pRLP44:RLP44-GFP (*bri1*^*cnu4*^) to depletion (PPZ) and exogenous supply of brassinosteroids. Bars indicate average relative root length ± S.D. (n =17 - 35).

**Supplemental Table S1**. Oligonucleotides used in this study

**Supplemental Table S2**. GreenGate Cloning modules and destination constructs

