## Supplemental Material for "A novel mutant allele uncouples brassinosteroid-dependent and independent functions of BRI1"

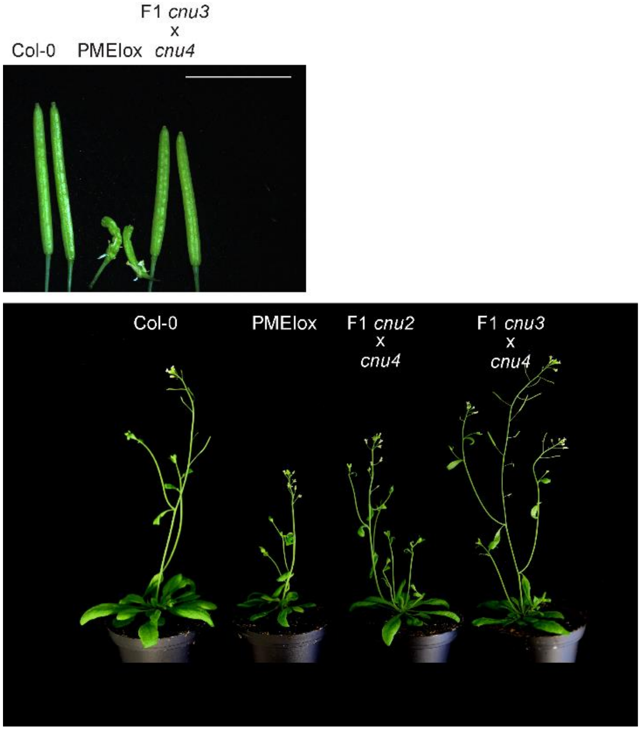


**Supplemental Figure 1.** *cnu3* and *cnu4* are allelic mutants. PMEIox silique morphology (upper panel) and plant stature (lower panel) remain suppressed in F1 plants of a cross between *cnu3* and *cnu4*, whereas F1 plants of a cross between *cnu2* (carrying a mutation in RLP44) and *cnu4* show PMEIox phenotype.

A

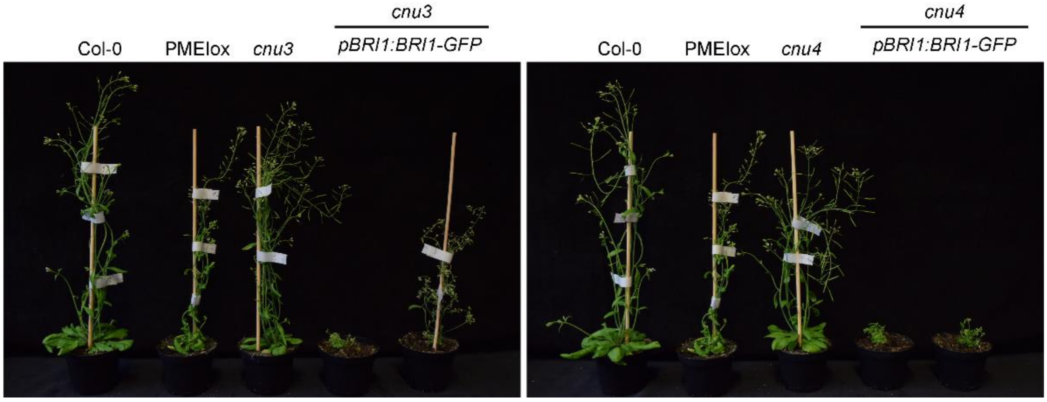


**Supplemental Figure 2.** Mutant BRI1 constructs complement the *cnu3* and *cnu4* mutants.

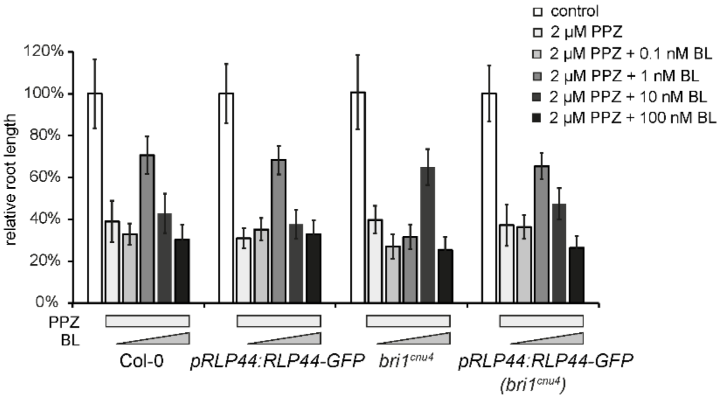


**Supplemental Figure 3.** RLP44 promotes BR response in the *bri1*$^{cnu4}$ mutant. Response of Col-0, pRLP44:RLP44-GFP, *bri1*$^{cnu4}$, and pRLP44:RLP44-GFP (*bri1*$^{cnu4}$) to depletion (PPZ) and exogenous supply of brassinosteroids. Bars indicate average relative root length ± S.D. (17 < n < 35).

**Supplemental Table S1**. Oligonucleotides used in this study

| oligonucleotide | sequence 5'-3' | remark |
|---|---|---|
| bri1cnu3_CAPS_F | TCGATTCCTGATGAGGTAGGTG | Cfr42I |
| bri1cnu3_CAPS_R | AAGATCCGCAAACGTGAGCTTC | Cfr42I |
| bri1cnu4_CAPS_F | TCAGGAGCTCATGTATGTCA | BseL1 |
| bri1cnu4_CAPS_R | TCCAATTGGTGTTGTTAGCAG | BseL1 |
| BRI1_attB1_L | GGGGACAAGTTTGTACAAAAAAGCAGGCTATGAAGACTTTTTCAAGCTTCTT | |
| BRI1_attB2_R | GGGGACCACTTTGTACAAGAAAGCTGGGTtTAATTTTCCTTCAGGAACTTCTT | |
| BRI1_GGC_1F | AACAGGTCTCAGGCTCATGAAGACTTTTTCAAGCTTCT | |
| BRI1_GGC_1R | AACAGGTCTCaATCACACGCGCCGGAGAGAAAGTCAG | |
| BRI1_GGC_2F | aacaGGTCTCaTGATACACTCACTGGaCTCGATCTCTCTGGA | |
| BRI1_GGC_2R | aacaGGTCTCaGAGtCCAGGATTGTTCAAGAA | |
| BRI1_GGC_3F | aacaGGTCTCaACTCTGTGGTTATCCTCTT | |
| BRI1_GGC_3R | aacaggTCTCaGGCCTCCTTCCATGAGATCT | |
| BRI1_GGC_4F | aacaGGTCTCaGgCCAGCGTCCCTTGCTGGT | |
| BRI1_GGC_4R | AACAGGTCTCACTGATAATTTTCCTTCAGGAACTTC | |
| at5G46630_cod_F | TCGATTGCTTGGTTTGGAAGAT | |
| at5G46630_cod_R | GCACTTAGCGTGGACTCTGTTTGC | |
| DWF4_F | caacagcaaaacaacggagcg | |
| DWF4_R | Tctgaaccagcacatagccttg | |

**Supplemental Table S2**. GreenGate Cloning modules and destination constructs

| | | |
|---|---|---|
| **pSW388** | **pBRI1:BRI1** | |
| pSW379 | BRI1(AT4G39400) promoter | Holzwart et al., 2018 |
| GGB003 | B-Dummy | Lampropoulos et al., 2013 |
| pSW380 | BRI1(AT4G39400) CDS | Holzwart et al., 2018 |
| pGGD002 | D-Dummy | Lampropoulos et al., 2013 |
| pGGE009 | UBQ10 terminator | Lampropoulos et al., 2013 |
| pGGF001 | pMAS::BastaR::tMAS | Lampropoulos et al., 2013 |
| pGGZ0001 | destination vector | Lampropoulos et al., 2013 |
| **pSW389** | **pBRI1:BRI1cnu3** | |
| pSW379 | BRI1(AT4G39400) promoter | Holzwart et al., 2018 |
| pGGB003 | B-Dummy | Lampropoulos et al., 2013 |
| pSW391 | BRI1cnu3 | This study |
| pGGD002 | D-Dummy | Lampropoulos et al., 2013 |
| pGGE009 | UBQ10 terminator | Lampropoulos et al., 2013 |
| pGGF001 | pMAS::BastaR::tMAS | Lampropoulos et al., 2013 |
| pGGZ001 | destination vector | Lampropoulos et al., 2013 |
| **pSW390** | **pBRI1:BRI1cnu4** | |
| pSW379 | BRI1(AT4G39400) promoter | Holzwart et al., 2018 |
| pGGB003 | B-Dummy | Lampropoulos et al., 2013 |
| pSW381 | BRI1cnu4 CDS | This study |
| pGGD002 | D-Dummy | Lampropoulos et al., 2013 |
| pGGE009 | UBQ10 terminator | Lampropoulos et al., 2013 |
| pGGF001 | pMAS::BastaR::tMAS | Lampropoulos et al., 2013 |
| pGGZ001 | destination vector | Lampropoulos et al., 2013 |
| **pSW427** | **pBRI1:BRI1cnu3,4** | |
| pSW379 | BRI1(AT4G39400) promoter | Holzwart et al., 2018 |
| pGGB003 | B-Dummy | Lampropoulos et al., 2013 |
| pSW419 | BRI1cnu3,4 | This study |
| pGGD002 | D-Dummy | Lampropoulos et al., 2013 |
| pGGE009 | UBQ10 terminator | Lampropoulos et al., 2013 |
| pGGF001 | pMAS::BastaR::tMAS | Lampropoulos et al., 2013 |
| pGGZ001 | destination vector | Lampropoulos et al., 2013 |
| **pSW421** | **pBRI1:BRI1cnu3-GFP** | |
| pSW379 | BRI1(AT4G39400) promoter | Holzwart et al., 2018 |
| pGGB003 | B-Dummy | Lampropoulos et al., 2013 |
| pSW391 | BRI1cnu3 | This study |
| pGGD001 | GFP | Lampropoulos et al., 2013 |
| pGGE009 | UBQ10 terminator | Lampropoulos et al., 2013 |
| pGGF001 | pMAS::BastaR::tMAS | Lampropoulos et al., 2013 |
| pGGZ001 | destination vector | Lampropoulos et al., 2013 |
| **pSW422** | **pBRI1:BRI1cnu4-GFP** | |
| pSW379 | BRI1(AT4G39400) promoter | Holzwart et al., 2018 |
| pGGB003 | B-Dummy | Lampropoulos et al., 2013 |
| pSW381 | BRI1cnu4 | This study |
| pGGD001 | GFP | Lampropoulos et al., 2013 |
| pGGE009 | UBQ10 terminator | Lampropoulos et al., 2013 |
| pGGF001 | pMAS::BastaR::tMAS | Lampropoulos et al., 2013 |

| | | |
|---|---|---|
| pGGZ001 | destination vector | Lampropoulos et al., 2013 |
| **pSW423** | **pBRI1:BRI1cnu3,4-GFP** | |
| pSW379 | BRI1(AT4G39400) promoter | Holzwart et al., 2018 |
| pGGB003 | B-Dummy | Lampropoulos et al., 2013 |
| pSW419 | BRI1cnu3,4 | This study |
| pGGD001 | GFP | Lampropoulos et al., 2013 |
| pGGE009 | UBQ10 terminator | Lampropoulos et al., 2013 |
| pGGF001 | pMAS::BastaR::tMAS | Lampropoulos et al., 2013 |
| pGGZ001 | destination vector | Lampropoulos et al., 2013 |
